# Tofacitinib overrides first-line drug resistance in a pre-clinical model of epilepsy

**DOI:** 10.64898/2026.08.06.743311

**Authors:** Jennifer Koehler, Olivia R. Hoffman, Quinn Rose Harvey, Barry A. Schoenike, Jose Ezekiel Clemente Espina, Avtar S. Roopra

## Abstract

One-third of people with epilepsy continue to have seizures despite antiseizure medications (ASMs), and available therapies often fail to improve disabling cognitive comorbidities. Patients with drug resistant epilepsy report that the adverse effects of medications along with their comorbidities can have a greater negative impact on the quality of life than seizures. We previously identified recurrent JAK/STAT3 activation in chronic epilepsy and showed that transient treatment with the JAK inhibitor tofacitinib (CP690550) durably suppresses seizures and restores cognition in mice. Here, we tested CP690550 as an add-on therapy after failure of carbamazepine (CBZ), a common first line treatment for epilepsy, in a mouse model of multifocal temporal lobe epilepsy. In CBZ-resistant animals, dual therapy with CP690550 reduced median seizure frequency and time spent seizing by an order of magnitude; most dual therapy responders had no observed behavioral seizures during treatment. CP690550 also restored spatial working and short-term memory. We found that cognitive rescue was independent of seizure response. Our work suggests that JAK/STAT inhibition can overcome ASM nonresponse while independently improving epilepsy-associated cognitive dysfunction.

## Introduction

Epilepsy is the fourth most common neurological disease, effecting over 65 million people worldwide*(1)*. Despite the FDA approval of 17 anti-seizure medications since 1975, 30% of patients have drug refractory disease*(2)*. Drug refractory epilepsy is defined as continuing seizures after treatment with two or more appropriate and well tolerated anti-seizure medications (ASM) *(3–7)*. However, only 10-15% of patients who fail to respond to one ASM will respond to the next treatment, and failure to respond to the first line of treatment is associated with overall drug resistance*(5, 7–11)*. Patients with drug resistant epilepsy overwhelmingly represent the greatest economic and psychosocial burden of the disease*(12)* as well as a greater burden of cognitive and behavioral comorbidities*(12–15)*. These comorbidities are often exacerbated by ASMs which have cognitive adverse effects of their own*(2, 16–18)*. Thus, there is a need to generate more efficacious treatments for patients who fail first lines of treatment and those who are resistant overall.

In acquired epilepsies, epileptogenesis represents a series of processes post an initial insult, such as a series of unremitting seizures known as status epilepticus (SE), that eventually results in spontaneous recurring seizures. Inflammatory signal cascades and astrogliosis have been shown to play a role in epileptogenesis*(19–24)*. Recent data has emerged both in patients*(25, 26)* and rodent models*(27, 28)* indicating that neuroinflammation could also play a role in drug resistant epilepsy. Inflammation has also been tied to cognitive decline in other neurological disorders including Alzheimer’s*(29, 30)*, Huntington’s disease*(31, 32)*, and age-related cognitive decline*(33–35)*. Our lab*(24, 36)* and others*(23, 37–39)* have identified the inflammatory Janus Associated Kinase/Singal Transductor and Activator of Transcription 3 (JAK/STAT3) as a driver of disease animal models and in patients with drug-refractory temporal lobe epilepsy.

We recently demonstrated that JAK/STAT3 signaling in chronic epilepsy provides a therapeutic target that when transiently targeted with CP690550, a pan JAK inhibitor with preference for JAK1*(40, 41)*, enduringly suppresses seizures and rescues cognitive decline*(24)*. Here we show that in epileptic mice that fail to respond to a first line of treatment, carbamazepine (CBZ), CP690550 is an effective add-on therapy. CP690550 reduced seizure frequency by 14-fold with a 70% response rate. Most responders had no observed behavioral seizures during treatment. Further, CP690550 restored spatial working and short-term memory. Finally, we provide evidence that seizure suppression is uncoupled from cognitive rescue. Thus, JAK/STAT inhibition can overcome ASM nonresponse while independently improving epilepsy-associated cognitive dysfunction.

## Methods

### Systemic kainic acid model

FVB/Nhsd mice were weighed and singly housed in observation chambers for the duration of the injections. Mice were injected intraperitoneally (i.p.) with synthetic kainic acid (KA) (7.5mg/kg for FVB) dissolved in 0.9% saline (#7065, Tocris Bioscience, Bristol, United Kingdom). At twenty-minute intervals, mice were given 7.5 mg/kg injections of KA up to the third injection; the dosage was then reduced to 5.0mg/kg. Animals continued to receive 5.0 mg/kg KA every twenty minutes for up to 10 injections. If an animal experienced two or more Class V or VI seizures within a single twenty-minute cycle, the subsequent KA injection was skipped. Injections resumed for the next round unless the animal reached status epilepticus (SE), and animals were considered to have reached SE after experiencing at least five Class V or VI seizures within a 90-minute window. During induction of SE, seizures were scored using a modified Racine Scale (86) where I = freezing, behavioral arrest, staring spells; II = head nodding and facial twitches; III = forelimb clonus, whole-body jerks or twitches; IV = rearing; V = rearing and falling; VI = violent running or jumping behavior. KA mice were observed for 1-2 hours after SE was achieved. Animals were returned to their home cages post SE. In the days following injection, animals were weighed and injected with 0.9% saline (s.c.) if body weight decreased by more than 0.5g and given gel diet (cat #) to aid in recovery.

### Generating a model of drug resistance

Following status epilepticus, a cohort of 16 mice were treated with carbamazepine (CBZ) for two weeks. CBZ (carbamazepine) (#4098, Tocris Bioscience, Bristol, United Kingdom) was dissolved in a solution of 30% PEG 400 (#39927-08-7, SIGMA, Darmstadt, Germany) in saline (0.9%). CBZ was administered via i.p. injections at a dose of 35 mg/kg twice daily 4 hours apart for two weeks in chronically epileptic mice, 12-14 weeks post SE. CP690550 (tofacitinib citrate) (#4556, Tocris Bioscience, Bristol, United Kingdom) (15mg/kg daily) was added on top of CBZ treatment for an additional two weeks to all CBZ treated mice (dual therapy). CP690550 was dissolved in a solution of 15% DMSO and 15% 100% EtOH in saline and administered via i.p. injections. All epileptic mice were handled daily and injected with vehicle to control for stress and handling. Treatment response defined as a 50% or greater reduction in seizure frequency, per clinical benchmarks*(42)*, as assessed via behavioral seizure recordings. CBZ treated mice were compared to baseline seizure frequency and dual therapy treated mice were compared to their CBZ treatment seizure frequency.

### Behavioral Seizure recording

To verify that mice had developed epilepsy, at 9 weeks post SE mice were video recorded for behavioral seizures for three weeks of baseline seizure recording followed by recording throughout treatment. Male and female FVB mice were singly housed in observation chambers for the duration of recording with access to food and water, and mice that did not exhibit at least 1 seizure per week (0.2/day) during the baseline recording period (or who died during recording) were triaged from the analysis. Videos were then reviewed and scored by a blinded experimenter using the modified Racine scale, and only class IV-VI were recorded.

### Y-maze cognitive testing

Working memory spontaneous alternation (SA) and short-term memory forced alternation (FA) tests were performed at 3 time points: prior to CBZ treatment (12 weeks post SE), post-CBZ treatment (14 weeks post-SE), and post dual therapy treatment (16 weeks post-SE). Historical data from naïve mice was used to compare for deficits. We also tested age-matched naïve mice treated with the dual therapy to test wither the compounds alone had cognitive adverse effects. All tests were video recorded and analyzed by an experimenter blinded to treatment. Mice were habituated in the testing room for 30 minutes prior to all trials. We used spontaneous alternation to evaluate working memory and forced alternation to evaluate short-term memory; both tests occurred within a three-arm Y-maze.

### Spontaneous Alternation

For spontaneous alternation, mice were placed in the starting arm of the maze and allowed to explore freely for 8 minutes. Sequential arm choices during exploration of a three-arm Y-maze are considered an indicator of spatial working memory because the innate curiosity of a healthy mouse promotes exploration of the arm least recently visited (36-38). The number of choices resulting in a spontaneous alternation (for example, an entry sequence of ABC but not ACA) was expressed as a percentage of the total number of choices. After prolonged time in the Y-maze, habituation to novelty, and thus changes in motivation, can reduce the percentage of spontaneous alternations over time*(43)*. Testing naïve mice, we observed the transition from an exploratory phase to an escape phase, and the total number of alternations for each mouse ranged from 20-100. To mitigate the effects of habituation and the variance in total alternation, we analyzed only the first 15 alternations of each mouse. Mice that did not reach 15 alternations were removed from the dataset. Distance traveled during each 8-minute test was measured to ensure that deficits in SA post-SE were not due to mobility impediments. Any mice who escaped the maze in the first 15 alternations were returned to the arm they escaped from, and the alternation count would start over. The maze was cleaned with 70% EtOH in between each trial. Mice that exhibited racing behavior, only moving in one circular motion for the entire 8 minutes were also excluded. All analysis was performed by an experimenter blinded to treatment conditions.

### Forced Alternation

Forced alternation (FA) tests depend on a mouse’s ability to recall and apply spatial short-term memory, motivated by their innate curiosity. FA tests involved two 5-minute trials 30 minutes apart: a sample trial and a retrieval trial. In the sample trial, the mouse was placed in the starting arm and allowed to explore the maze with one arm blocked off. Between trials and mice the maze was washed with 70% EtOH. In the retrieval trial, the previously blocked arm was opened, and the mouse was allowed to explore freely for 5 minutes. Mice that did not enter all three arms of the maze during the retrieval trial of FA tests were excluded from the dataset. Mice that exhibited racing behavior, only moving in one circular motion for the entire 8 minutes were also excluded. FA tests were quantified as percentage of time spent in the novel arm. A mouse was considered inside the novel arm when its hindlimbs entered it, and outside the novel arm when its hindlimbs exited. Time spent in the center of the maze was not counted. All analysis was performed by an experimenter blinded to treatment conditions.

### Tissue isolation and homogenization for western blot

After KA seizure induction or treatment with the dual therapy, animals were sacrificed by decapitation. Whole hippocampal hemispheres were harvested and flash-frozen in liquid nitrogen or on dry ice. Hippocampal tissue was lysed in Radioimmunoprecipitation Assay Buffer (RIPA: 50mM Tris, 150mM NaCl, 1% nonidet P-40, 0.5% sodium deoxycholate, 0.1% SDS) with mammalian protease inhibitor (1:1000, Sigma or 1860932 ThermoScientific) and phosphatase inhibitor (1:1000, Sigma or 78428 ThermoScientific). Protein from cell lines was harvested in Triton lysis buffer (3% 5M NaCl, 10% glycerol, 0.3% Triton X-100, 5% 1M Tris pH8.0) after PBS washes. Tissue was homogenized in lysis buffer by probe sonication (Fisher Scientific, Sonic Dismembrator, Model 100, Hampton, NH) on power 3 for two rounds, with 10 pulses per round. Supernatants were isolated by centrifugation and quantified using the DC Protein Assay (Bio-Rad, Hercules, CA) or BCA assay (ThermoScientific 23227). Protein extracts were stored at -80°C. 5x loading buffer (0.5mM Tris, 10% SDS, 50% glycerol, 10mM EDTA, and 1% beta-mercaptoethanol) was added to each sample to reach a 1x final concentration. Extracts in loading buffer were boiled at 95°C for 5 minutes and stored for up to one month at -20°C until run on an acrylamide gel.

### Western Blotting

Following systemic kainite induced epilepsy, protein extracts in loading buffer were loaded at 20μg per lane and resolved by electrophoresis in hand-poured acrylamide gels with a 5% acrylamide stacking layer (125mM Tris pH6.8, 5% acrylamide, 0.01% ammonium persulfate, 0.01% SDS, 0.01% TEMED) and an 8% acrylamide separating layer (375mM Tris pH8.8, 8% acrylamide, 0.015% ammonium persulfate, 0.015% SDS, 0.08% TEMED). Gels were transferred to polyvinyl difluoride membranes (PVDF; Millipore, Bedford, MA) using Tris-glycine transfer buffer (20mM Tris, 1.5M glycine, 20% methanol). Membranes were blocked with 5% bovine serum albumin (BSA) (Fisher Scientific, Fair Lawn, NJ) diluted in low-salt Tris-buffered saline (w/w TBST; 20mM Tris pH7.5, 150mM NaCl, 0.1% Tween 20) with mammalian protease inhibitor (1:2000) and phosphatase inhibitor (1:2000), for 1 hour at room temperature. Primary antibodies were diluted in the same blocking buffer and incubated with membranes overnight at 4°C. Antibodies include: phospho-STAT3 (Tyr705) (1:2000, #9145 Cell Signaling), total STAT3 (1:1000, #12640 Cell Signaling), Tubulin (1:1000, #T9026 Sigma-Aldrich). The next day, the membranes were washed three times in 1X TBST and incubated with horseradish peroxidase-conjugated goat-anti-mouse or -rabbit secondary antibodies for one hour at room temperature (1:10,000, Invitrogen #31430, #31460, Rockford, IL). Membranes were subsequently washed three times in TBST, and membranes were developed in SuperSignal West Femto (pSTAT3) or SuperSignal West Pico (total Stat3, Tubulin) reagent (ThermoFisher, Waltham, MA). Bands were imaged using a ChemiDoc-It Imaging System (UVP Vision-Works, Upland, CA) and quantified using UVP Vision Works software or ImageJ. Band intensities were graphed and analyzed using Prism 9 software (GraphPad Software, La Jolla, CA).

### Flurothyl seizure threshold testing

All seizure threshold tests were video recorded, and seizure behavior was scored by a blinded observer. Flurothyl seizure threshold tests were CBZ treated mice and vehicle control mice using 100% bis(2,2,2-trifluoroethyl) ether (#287571, Sigma-Aldrich, St. Louis, MO). All tests were performed in a fume hood. Mice were placed in an airtight 10L. Plexiglas chamber (8.5”X10.75”X6.75”). Fluorothyl was infused into the chamber using a peristaltic pump at a rate of 40μL/minute onto a piece of Whatman filter paper placed at the top of the chamber. The time to generalized tonic clonic seizure (GTCS) was recorded from the start of fluorothyl infusion. Mice reached GTCS when they exhibited a complete loss of postural/motor control. After experiencing a generalized tonic clonic seizure, mice were rapidly removed from the chamber, and fluorothyl infusion was terminated. Mice were sacrificed and brains were harvested immediately post seizure threshold testing.

### Brain and blood collection for pharmacokinetics

Brain hemispheres were collected from 5 CBZ treated naive animals this occurred immediately post flurothyl testing (1, 4, and 8 hours post i.p. injection). The hemispheres were stored at -80C until analysis.

For CP690550 pharmacokinetics analysis serum and brain was collected from 4 chronically epileptic mice 15 minutes after the tenth injection of CP690550 (15mg/kg) and 1 hour after a single injection of trospium chloride (1 mg/kg in saline, i.p.). Serum was collected via cardiac puncture into 0.5M EDTA coated syringes and transferred into 0.5 ml Eppendorf tubes. Plasma was isolated via centrifuge (2000 rpm for 10 minutes at 4 degree C) and stored at -80 degrees C. The brain hemispheres were rapidly collected after cardiac puncture, separated into hemispheres and stored at -80 degrees C.

### Bioanalysis

#### Materials

Bioanalysis was performed by the University of Michigan Core labs as follows. Tofacitinib was obtained from Tocris (#4556, Tocris Bioscience, Bristol, United Kingdom). Trospium chloride reference standards were purchased from Tocris (#T3305, Tokyo Chemical Industry Co. Ltd). Carbamazepine reference standard was purchased from (#4098, Tocris Bioscience, Bristol, United Kingdom). IPI549 (HY-100716, Eganelisib; MedChemExpress) was used as the internal standard (IS). Acetonitrile (LC-MS grade) was obtained from MilliporeSigma (AX0156-1, OmniSolv, Supelco). Formic acid (99.0+%) (A117-05AMP, Optima™ LC/MS grade) was purchased from Fisher Chemical™ (Fisher Scientific). Dimethyl sulfoxide (DMSO, HPLC grade, product no. 34869) was purchased from Sigma-Aldrich.

### Standard

A primary stock solution of was prepared in DMSO at 5mg/ml for CBZ, and 10mg/ml for tofacitinib and trospium chloride. Serial dilutions in acetonitrile yielded working standard solutions at 5000, 2500, 1000, 500, 250, 100, 50, 25, 10, 5, 2.5, and 1 ng/mL for CBZ with brain homogenate. Calibration standards were prepared in blank brain homogenate by mixing 30 µL of each working standard solution with 30 µL of blank brain homogenate and 100 µL of IS solution Quality control (QC) stock solutions were prepared independently from a separate weighing and diluted to four concentration levels (5, 250, 1000, and 2500 ng/mL) in acetonitrile.

Additional serial dilution generated working standards for CP690550 and TCL at 1000, 500, 250, 100, 50, 25, 10, 5, 2.5, and 1ng/mL for serum and 100, 50, 25, 10, 5, 2.5, and 1 ng/mL for brain homogenate. Quality control (QC) stock solutions were prepared independently from a separate weighing and diluted to four concentration levels (5, 250, 1000, and 2500 ng/mL) in acetonitrile.

Calibration curves were constructed using at least 6 non-zero standards by plotting the peak area ratio of each analyte to IS against nominal concentration, fitted by linear regression with 1/x² weighting. Linearity was confirmed by correlation coefficients r > 0.99 for both analytes in both matrices.

### Brain Tissue Homogenization

Brain tissues were homogenized at a 1:5 (w/v) ratio in 20% acetonitrile/water (v/v) using a 2 mL soft tissue homogenizing kit (Precellys Lysing Kit, cat. no. P000918-LYSK1, Bertin Technologies) on a Precellys Evolution homogenizer equipped with a CryoLys Evolution cooling system (Bertin Technologies). Homogenization was performed at 10°C, with 2 runs of 4 cycles each at 6,800 rpm for 20 seconds per cycle.

### Brain Homogenate Sample Preparation

Calibration standards in brain homogenate were prepared by mixing 50 µL of each working standard solution with 50 µL of blank brain homogenate and 50 µL of IS solution (IPI549, 50 ng/mL in acetonitrile) in a 96-well plate. Study samples were prepared by combining 50 µL of brain homogenate with 50 µL of acetonitrile and 50 µL of IS solution. All samples were vortexed for 10 minutes and centrifuged at 4,000 rpm for 10 minutes. The resulting supernatant was transferred to a new 96-well plate, and 5 µL was injected for LC-MS/MS analysis.

### Serum Sample Preparation

Calibration standards in mouse serum were prepared by mixing 40 µL of each working standard solution with 40 µL of blank serum and 100 µL of IS solution (IPI549, 50 ng/mL in acetonitrile) in a 96-well plate. Study samples were prepared by combining 40 µL of serum with 40 µL of acetonitrile and 100 µL of IS solution. All samples were vortexed for 10 minutes and centrifuged at 4,000 rpm for 10 minutes. The resulting supernatant was transferred to a new 96-well plate, and 5 µL was injected for LC-MS/MS analysis.

### LC-MS/MS Analysis

Quantitative analysis for CP690550 and TCL was performed on an AB Sciex 5500 triple quadrupole mass spectrometer equipped with a TurboIonSpray® source (AB Sciex LLC, Redwood City, CA, USA) operated in positive ion mode. Chromatographic separation was performed on a Waters XBridge C18 column (50 mm × 2.1 mm, 5.0 µm particle size) using a binary gradient of 0.1% formic acid in water (mobile phase A) and 0.1% formic acid in acetonitrile (mobile phase B) at a flow rate of 0.5 mL/min. The gradient program was as follows: held at 5% B from 0.01 to 0.50 min, ramped linearly to 98% B at 1.00 min, held at 98% B until 2.80 min, returned to 5% B at 2.90 min, and re-equilibrated until 5.00 min.

The quantitation analysis for CBZ was performed on an AB Sciex 4500 triple quadrupole mass spectrometer equipped with a TurboIonSpray® source (AB Sciex LLC, Redwood City, CA, USA) operated in positive ion mode. Chromatographic separation was performed on a Waters XBridge C18 column (50 mm × 2.1 mm, 3.5 µm particle size) using a binary gradient of 0.1% formic acid in water (mobile phase A) and 0.1% formic acid in acetonitrile (mobile phase B) at a flow rate of 0.4 mL/min. The gradient program was as follows: held at 5% B from 0.01 to 0.50 min, ramped linearly to 95% B at 1.50 min, held at 95% B until 3.50 min, returned to 5% B at 3.60 min, and re-equilibrated until 5.60 min.

Mass spectrometric detection was performed in positive ionization mode using multiple reaction monitoring (MRM) with a 5 ms pause between transitions. Monitored transitions were *m/z* 237.1 → 193.9 for carbamazepine (DP 84 V, EP 11V, CE 5 V, CXP 17 V) and *m/z* 529.2 → 326.3 for IPI549 (DP 100 V, EP 10, CE 56.9 V, CXP 16.13 V). Monitored transitions were *m/z* 313.1 → 148.7 for tofacitinib (DP 114 V, EP 12.7 V, CE 42 V, CXP 9 V), *m/z* 393.2 → 182.3 for trospium chloride (DP 81 V, EP 6.7 V, CE 47 V, CXP 16 V), and *m/z* 529.2 → 337.1 for IPI549 (DP 100 V, EP 12 V, CE 30 V, CXP 16 V). Secondary confirmatory transitions were monitored at *m/z* 313.1 → 172.9 for tofacitinib and *m/z* 393.2 → 164.2 for trospium chloride.

For CP690550 and TCL the LLOQ was 2.5 ng/mL in serum and 1 ng/mL in brain homogenate, corresponding to 5 ng/g in brain tissue for both analytes. For CBZ the LLOQ was 2.5 ng/mL in brain homogenate, corresponding to 12.5 ng/g in brain tissue. Intra-batch accuracy ranged from 88.0% to 121.3% of nominal concentration, with precision (%RSD) ≤ 18.9% across all QC levels, meeting the pre-defined acceptance criteria of ≤ 15% deviation from nominal (≤ 20% at the LLOQ).

### Animal care

All animal procedures and experiments were performed with approval from the University of Wisconsin-Madison School of Medicine, Public Health Institutional Animal Care and Use Committee, and according to NIH national guidelines and policies.

### General

Male and female FVB mice were bred and housed under a 12-hour light/dark cycle with access to food and water ad libitum. Mice were allowed to reach 5-7 weeks of age (FVB) before undergoing experimentation. Kainic acid injections were performed at the same time of day approximately 9 A.M., and mice were returned to home cages before the start of the dark cycle (approximately 5 P.M.). Mice were housed with littermates, with males and females separated at weaning.

### Statistical analysis

For all statistical analysis—unless otherwise specified—*q* < 0.05 or *P* (corrected for multiple comparisons) <0.05 was considered statistically significant (\**P*/*q* < 0.05, \*\**P*/*q* < 0.01, \*\*\**P*/*q* < 10^−3^, and \*\*\*\**P*/*q* < 10^−4^). Data were graphed as means ± SD in Prism 10 software (GraphPad Software, La Jolla, CA). Statistical tests were also performed in Prism. Data were analyzed by *t* test, two-way ANOVA/mixed-effects models, or one-way ANOVA, with the two-stage step-up method of Benjamini, Krieger, and Yekutieli (abbreviated BKY in figure legends). Paired analysis was performed using the Friedman test with BKY post hoc for two or more conditions. Kruskal-Wallis tests with Benjamini, Krieger, Yekutieli correction for data with >2 conditions. Non-linear regression analysis was performed for pharmacokinetic data. Correlation analysis was performed using the Spearman test. All statistical tests were two-tailed.

## Results

### CBZ exhibits expected brain pharmacokinetics

To confirm that pharmacokinetic studies of carbamazepine (CBZ) in other mouse strains eg C57B6(REF)s carry over to FVB mice that are used herein, we first quantified brain exposure and anticonvulsant activity of CBZ following a single intraperitoneal injection (35 mg/kg). Brain concentrations of CBZ declined over the first 8 hours post-injection from 6606ng/g± 2117 to 50ng/g +/- 94 **(Fig. 1A).** This temporal profile was paralleled by a reduction in anticonvulsant efficacy, with seizure threshold significantly decreased between 4 and 8 hours post-injection (Kruskal–Wallis, q = 0.0384) (**Fig. 1B**). Consistent with a pharmacokinetic–pharmacodynamic relationship, CBZ brain concentration was positively associated with seizure threshold (nonlinear regression, R² = 0.7063, p = 0.0002) (**Fig. 1C**), confirming dependence of anticonvulsant efficacy on drug concentration.

**Fig 1.**
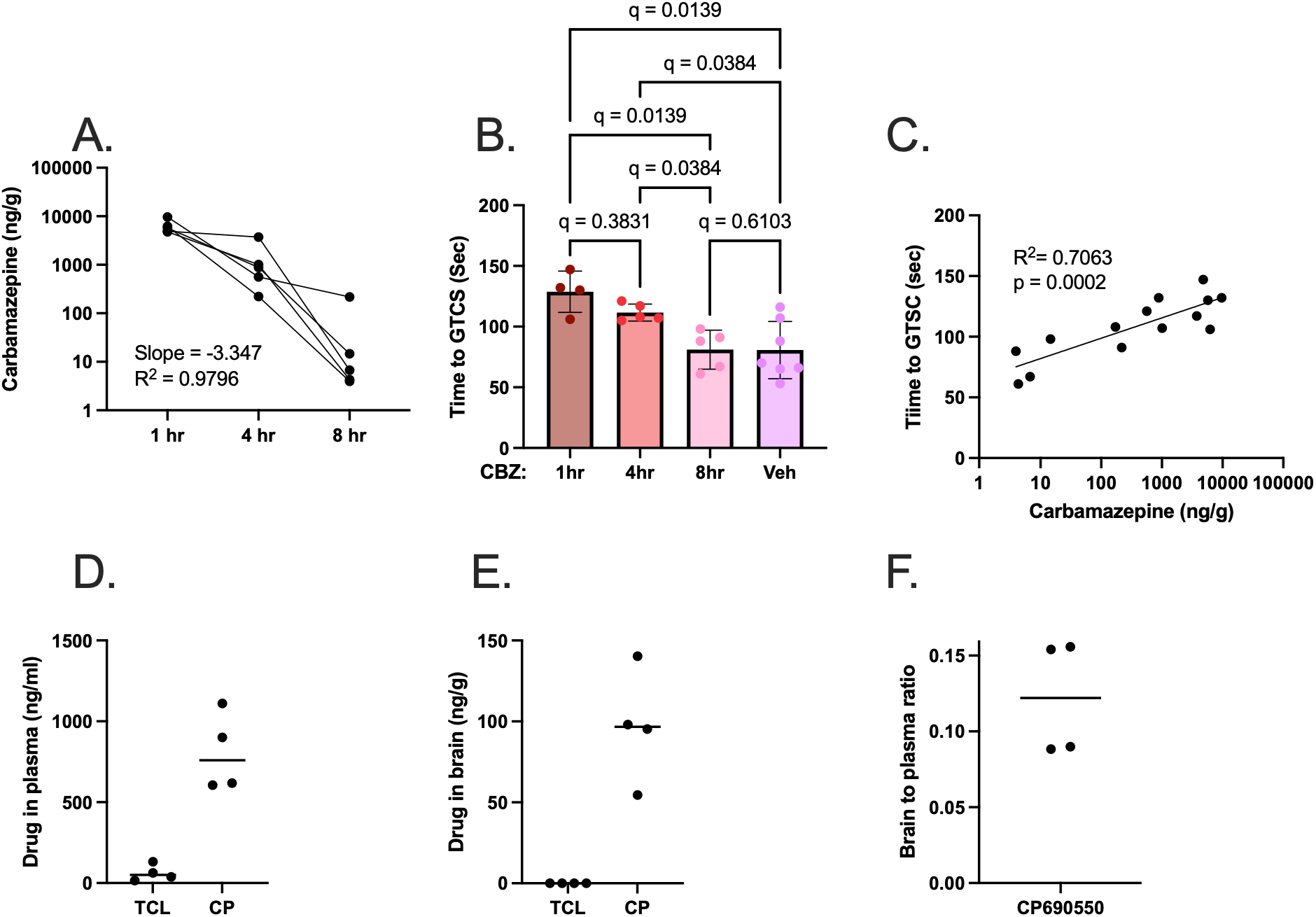
Both CBZ and CP690550 cross the blood brain barrier. **(A)** Brain concentrations (ng/g) of carbamazepine at 1 hour, 4 hours and 8 hours after a single injection. **(B)** Time to first general tonic clonic seizure after exposure to fluoroethyl in mice treated with one dose with CBZ 1 hour, 4 hour, and 8 hours after i.p. injection as well as a vehicle control Kruskal-Wallis test with BKY post hoc. **(C)** Correlation between seizure threshold and CBZ concentration in the brain. **(D)** Serum and **(E)** brain concentrations of Trospium Chloride (TCL), a non-blood brain barrier penetrant molecule, and C690550 (CP) in chronically epileptic mice are depicted along. **(F)** brain to plasma ratio of CP690550. Mean denoted by horizontal line

### CP690550 shows brain penetration in chronically epileptic mice

We assessed brain exposure of CP690550 in chronically epileptic mice, a condition in which blood–brain barrier integrity may be altered. The mean serum concentration was 808.5 ng/mL **(Fig 1D)**, and mean brain concentration was 96.9 ng/g (∼310.2 nM) **(Fig 1E)** resulting in a brain-to-plasma ratio of 0.12 **(Fig. 1F)**. In contrast, Trospium Chloride, a compound known not to penetrate the blood brain barrier, remained below the limit of detection. We note that the brain concentrations of CP690550 exceeds 89% of the reported in vitro whole blood IC₅₀ values for JAK1 heterodimers*(40)*.

### The repeated low dose systemic kainic acid mouse model is a model of CBZ resistance

We previously demonstrated the seizure suppressing effects of CP690550 during chronic epilepsy*(24)*. To test whether CP690550 would be effective in mice that experience persistent seizures despite conventional ASM treatment, we used carbamazepine (CBZ) to screen for drug-resistance prior to the addition of CP690550 (dual therapy). To do this, we induced status epilepticus (SE) with systemic kainic acid (KA) i.p. injections. Given the tight association of behavioral seizure detection by video and electrographic seizure detection by EEG in this model *(24)*, we used video monitoring for behavioral seizures. Video recording began 9 weeks post SE. Mice were treated with CBZ at 12 weeks post SE (35mg/kg twice daily 4 hours apart) for two weeks followed by the addition of CP690660 (15mg/kg, daily) on top of continued CBZ treatment **(Fig 2A)**. We screened mice for drug resistance using the clinical benchmark of a reduction in epileptic baseline seizure frequency of 50% or more. Consistent with other temporal lobe epilepsy mouse models *(44–47)* 38% of mice were responsive to CBZ whereas 62% failed to reach the clinical definition of ASM response **(Fig 2B).**

**Fig 2.**
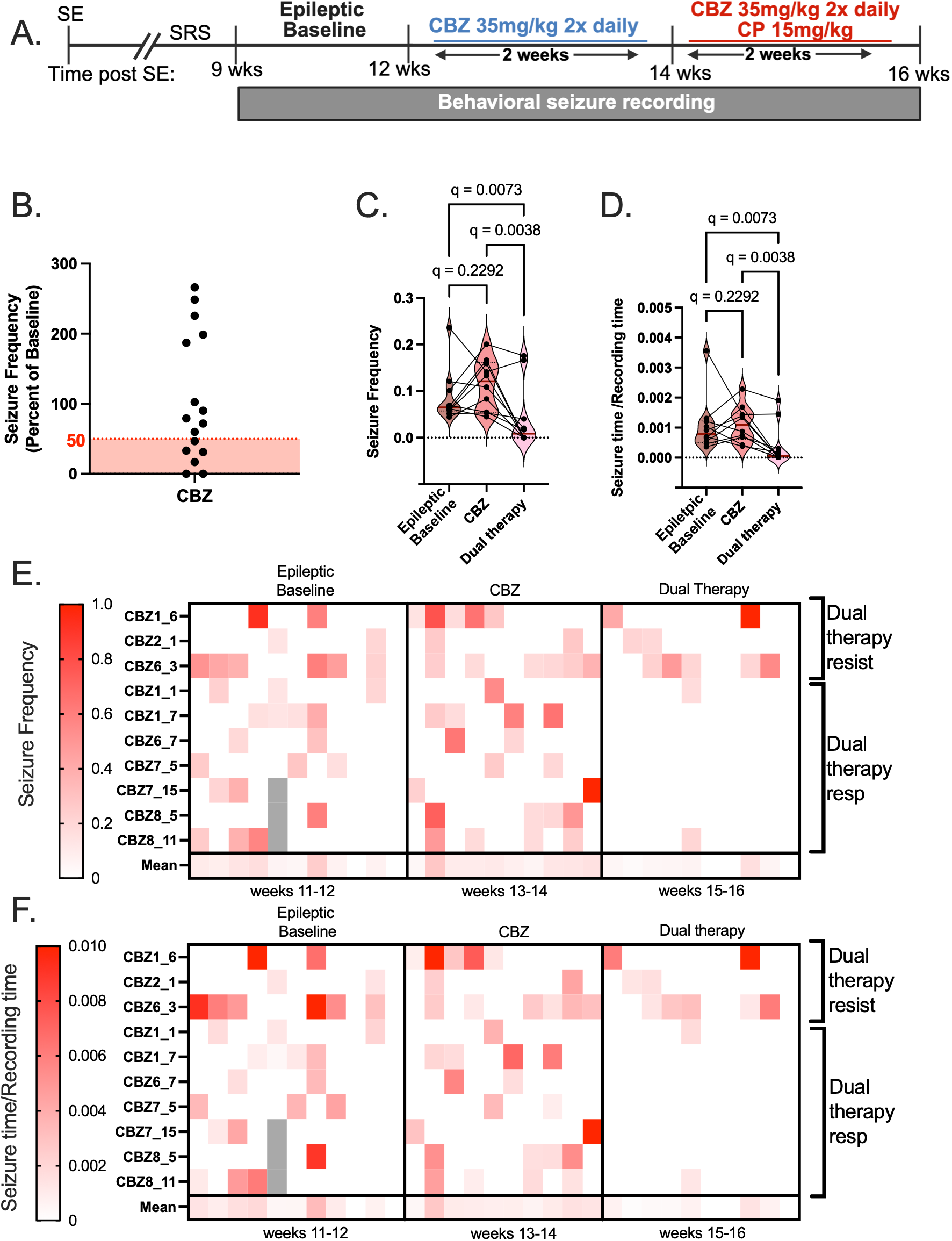
CP690550 suppresses seizure in CBZ resistant mice. **(A)** Experimental timeline**. (B)** Seizure frequency of CBZ treated mice is depicted as a percent of baseline. Mice with a reduction in seizure frequency greater than 50% (denoted by the red shaded area) are CBZ responder (n=6) and those above that threshold are non-responders (n=10). **(C)** Seizure frequency and **(D)** time spent seizing for all CBZ non-responders. The median is denoted by a bold maroon line. All statistics performed using a Friedman test with BKY post hoc. Heat maps depict individual mouse responses across epileptic baseline, CBZ treatment and dual therapy treatment for both **(E)** seizure frequency and **(F)** time spent seizing. Brackets denote dual therapy responders and non-responders on the heatmaps.

### CP690550 treatment suppresses seizures in CBZ resistant mice

We next assessed whether CP690550 could suppress seizures in the cohort of CBZ resistant. Dual therapy led to a 14-fold reduction in median seizure frequency and an 18-fold reduction in median time spent seizing compared to CBZ treatment alone (Friedman test, frequency: q = 0.0038, time spent seizing: q = 0.0038) **(Fig 2C-F).** Of the 62% of mice that were resistant to CBZ, 70% responded to dual therapy. In aggregate, combining the number of mice that originally responded to CBZ alone and those that responded to dual therapy, a total of 82% of mice had achieved seizure suppression greater than or equal to 50%.

### CP690550 dual therapy profoundly suppresses seizures in mice which respond to treatment

We noted 2 populations of mice upon CP690550 treatment: mice that did not respond at all to dual therapy and those that showed robust reductions in seizure frequency and burden **(Fig 2E-F).** To explore this further, we asked how well CP690550 suppressed seizures in those mice that showed a greater than 50% reduction in seizure frequency with dual therapy compared to CBZ alone. In the 70% of mice that showed greater than 50% reduction in seizure frequency with dual therapy compared to CBZ alone, the majority of mice were seizure free during the recording window and exhibited a 100-fold reduction in both median seizure frequency (Freidman test, q = 0.0034) and median time spent seizing (Friedman test, q = 0.0034) compared to CBZ treatment **(Fig 3B-C).** Five of 7 CBZ+CP690550 responders were free from behavioral seizures during video recordings while on dual therapy and the remaining 2 mice had only a single seizure each **(Fig 2D-E)**. We also see a 100-fold reduction in median seizure frequency (Friedman test, q = 0.0015) and time spent seizing (Friedman test, q = 0.0015) for all dual therapy responders compared to CBZ treatment, regardless of CBZ resistance status **(Fig S1-2).** Note that CBZ responders show only a 3-fold reduction in both median seizure frequency (Friedman test, q = 0.0160) and time spent seizing (Friedman test, q = 0.0074) **(Fig S3)**. These results suggest that mice which respond to dual therapy, do so robustly.

**Fig 3.**
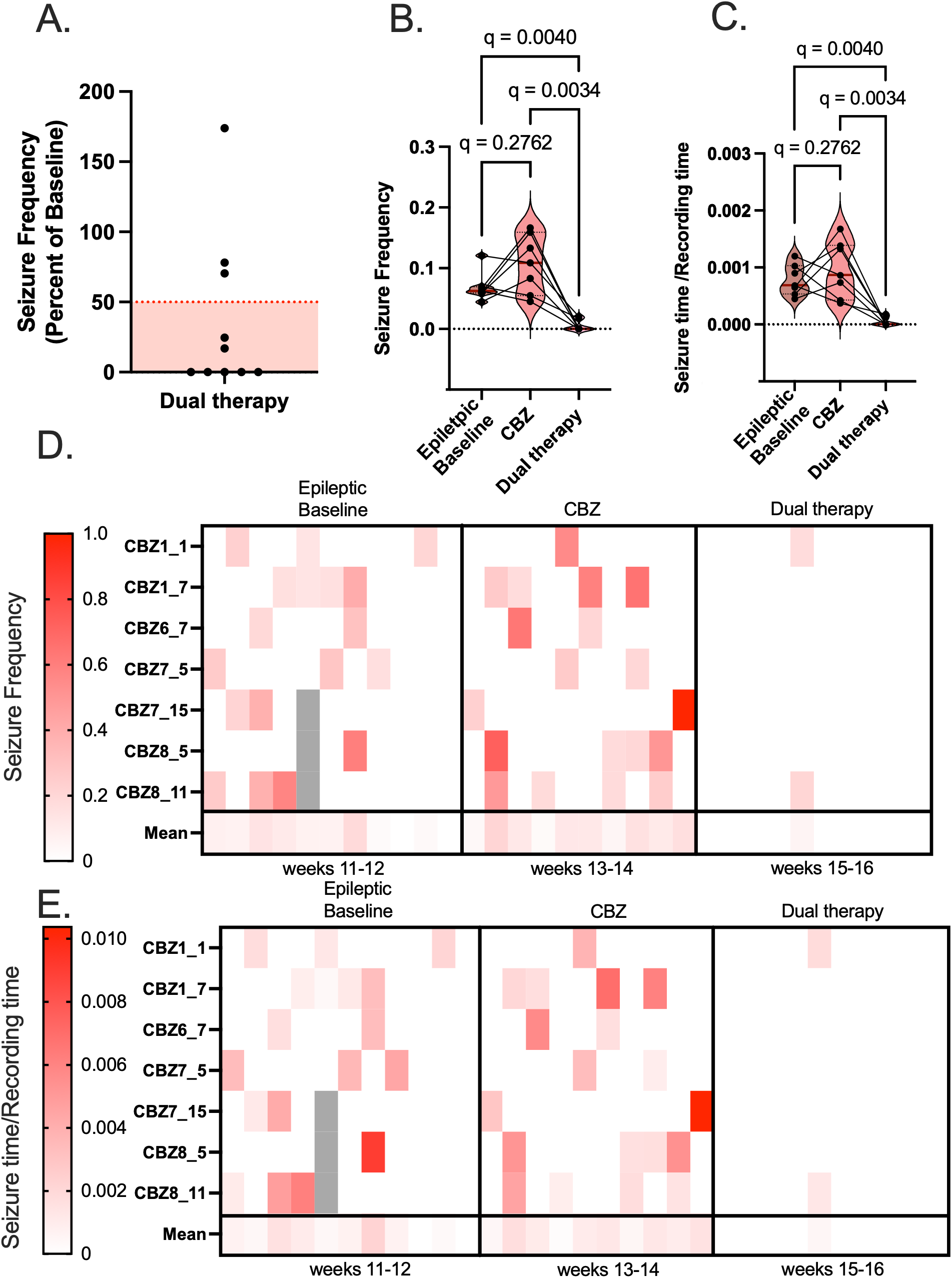
Most drug-resistant mice that respond to CP690550 dual therapy achieve seizure freedom. **(A)** The seizure frequency of dual therapy treated mice is depicted as a percent of epileptic baseline, mice with a reduction in seizure frequency (seizures/hour) greater than 50%, the red shaded area, are dual therapy responders, n = 7 and those above that threshold are non-responders, n = 3. Violin plots represent the **(B)** seizure frequency and **(C)** time spent seizing for all CBZ non-responders and dual therapy responders. The median is denoted by a bold maroon line. All statistics performed using a Friedman test with BKY post hoc. Heat maps depict individual mouse responses across epileptic baseline, CBZ treatment and dual therapy treatment for both **(D)** seizure frequency and **(E)** time spent seizing in drug resistant mice which respond to dual therapy treatment.

### CP690550 rescues spatial memory in a mouse model of drug-resistant epilepsy

We have previously shown that a transient two-week treatment with CP690550 in chronically epileptic mice rescued working and short-term memory with effects lasting at least two weeks post drug removal*(24)*. To test spatial memory, we utilized the spontaneous alternation y maze test and forced alternation Y maze test as previously described*(24)*. Both tests leverage the inherent curiosity of mice to test their spatial working and short-term memory respectively. The spontaneous alternation test allows mice to explore a Y maze for 8 minutes and quantifies the number of sequential alternations that occur during the first 15 alternations. This percent spontaneous alternation measures working memory since mice must recall the arm most recently explored to make a spontaneous alternation into the most novel arm. The forced alternation test measures short-term memory by quantifying the percentage of time mice spend in a novel arm for 5 minutes when the mice were pre-exposed to the other arms of the maze in a sample trial 30 minutes prior. To test whether CP690550 could rescue spatial memory deficits in CBZ resistant epilepsy, we performed cognitive testing prior to CBZ treatment (epileptic baseline), after 2 weeks of CBZ treatment (CBZ), and after two weeks of dual therapy treatment (dual therapy) **(Fig 4A).** We deployed the spontaneous and forced alternation y maze tests to quantify working and short-term memory respectively **(Fig 4B).** We show clear deficits in the percent spontaneous alternations for epileptic mice compared to naïve which CBZ fails to rescue (naïve vs epileptic Kruskal-Wallis q < 0.0001, epileptic vs CBZ Kruskal-Wallis q = 0.2300) **(Fig 4C)**. In contrast dual therapy for 2-weeks was associated with the restoration of working memory to levels comparable to naïve mice (CBZ vs dual therapy Kruskal-Wallis q = 0.0055, dual therapy vs naïve non-epileptic Kruskal-Wallis q = 0.2300) **(Fig 4C)**. CBZ treated mice also showed deficits in short-term memory compared to naïve mice (Kruskal-Wallis, q < 0.0001) **(Fig 4D)**. The addition of CP690550 rescues short-term memory up from both epileptic baseline and CBZ treatment levels and back to naïve levels (epileptic baseline vs dual therapy Kruskal-Wallis q = 0.0019, CBZ vs dual therapy Kruskal-Wallis q = 0.0002, dual therapy vs naïve non-epileptic Kruskal-Wallis q = 0.1284) **(Fig 4D)**. These data suggest that dual therapy treatment successfully restores spatial and working memory after a previous anti-seizure treatment has failed.

**Figure 4.**
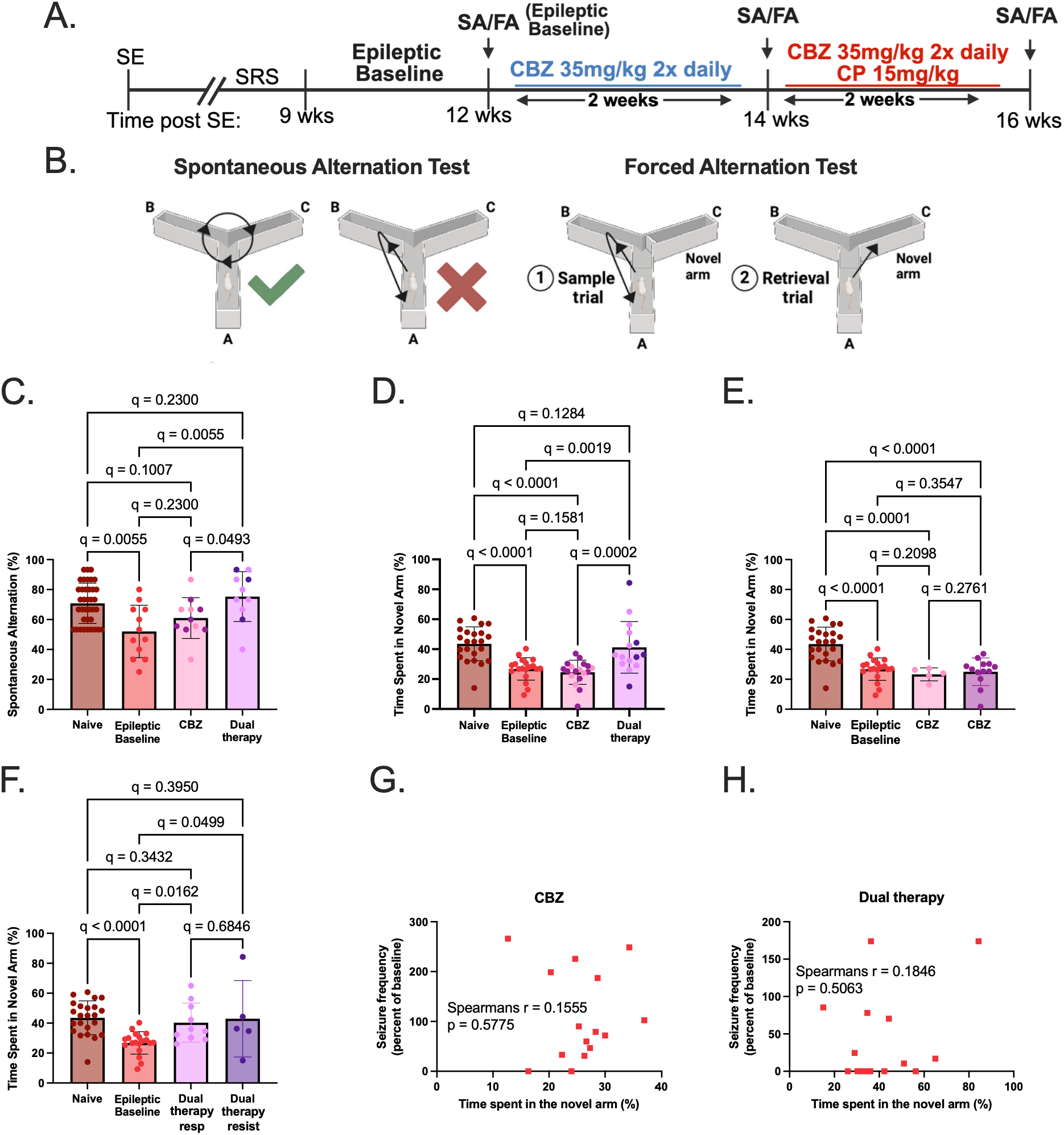
CP690550 restores working and short-term memory after CBZ treatment fails. **(A)** Experimental timeline. Working memory was quantified using the spontaneous alternation y maze test (SA) to measure the percentage of novel, or spontaneous, alternations during an 8-minute interval. Short-term memory was measured using the forced alternation y maze (FA) where the percent time spent in the novel arm was measured. **(B)** Visual representation of the spontaneous alternation test for working memory and the forced alternation test for short-term memory. We measured **(C)** working memory as percent spontaneous alternation and **(D)** short term memory as time spent in the novel arm for naïve, epileptic, CBZ, and dual therapy treated mice. Each dot represents a mouse; the darker dots in the CBZ and dual therapy bars represent mice who did not show seizure suppression. Spatial short-term memory was further quantified for seizure responders for both **(E)** CBZ and **(F)** dual therapy. All statistics were performed using the Kruskal-Wallis test with BKY post-hoc. Spearman’s correlation analysis of percent reduction in seizure frequency and short-term memory for both **(G)** CBZ and **(H)** dual therapy.

### Cognitive rescue is uncoupled from seizure suppression

To test whether cognitive rescue is correlated with seizure suppression we separated CBZ and dual therapy treated mice into responders and resistant groups based on seizure suppression using the clinical threshold described previously. We find that for CBZ treated mice, deficits in short-term memory persist both in mice that experienced a 50% reduction in seizure frequency and those that did not (naïve non-epileptic vs CBZ responders Kruskal Wallis q = 0.0001, naïve non-epileptic vs CBZ non-responders Kruskal-Wallis, q < 0.0001) **(Fig 4E)**. CP690550’s ability to restore cognition is also independent of seizure response, with dual therapy treated seizure-responders, and seizure non-responders exhibiting a significant rescue in short term memory compared to epileptic baseline (dual therapy responders vs epileptic baseline Kruskal-Wallis, q = 0.0162, dual therapy non-responders vs epileptic baseline Kruskal-Wallis, q = 0.0499) **(Fig 4F)**. We also see no significant correlation between fold change in seizure frequency and time spent in the novel arm for either CBZ (Spearman correlation, p = 0.5775) or dual therapy treatment (Spearman correlation, p = 0.5063) **(Fig 4G-H).** Together, these data provide evidence uncoupling cognitive rescue from seizure suppression.

### CP690550 dual therapy suppresses pSTAT3 in regardless of seizure response

To test this hypothesis that mice continue to have uncontrolled seizures when treated with dual therapy due to a lack of target engagement we measured STAT3 activation by western blot **(Fig 5A).** Previous work by our lab has shown that epileptic mice have increased pY705 in the hippocampus compared to age matched naïve mice in the chronic period*(24)*. We recapitulate this result (one-way ANOVA, q = 0.0003) while showing that treatment with dual therapy for two weeks significantly suppresses pY705-STAT3 compared to epileptic mice in both CBZ+CP690550 responders and dual therapy resistant mice (epileptic vs dual therapy responders one-way ANOVA, q < 0.0001, epileptic vs dual therapy non-responders one-way ANOVA q = 0.0003) **(Fig 5B)**. Interestingly, we find an increase in total STAT3 in our dual therapy treated mice compared to naïve mice (dual therapy responders vs naïve mice one-way ANOVA q = 0.0065, dual therapy non-responders vs naïve one-way ANOVA q = 0.0019) **(Fig 5C)**. Together these data indicate that a failure to suppress seizures present in a subset of dual therapy treated mice is not due to failure to suppress the STAT3 activation.

**Figure 5.**
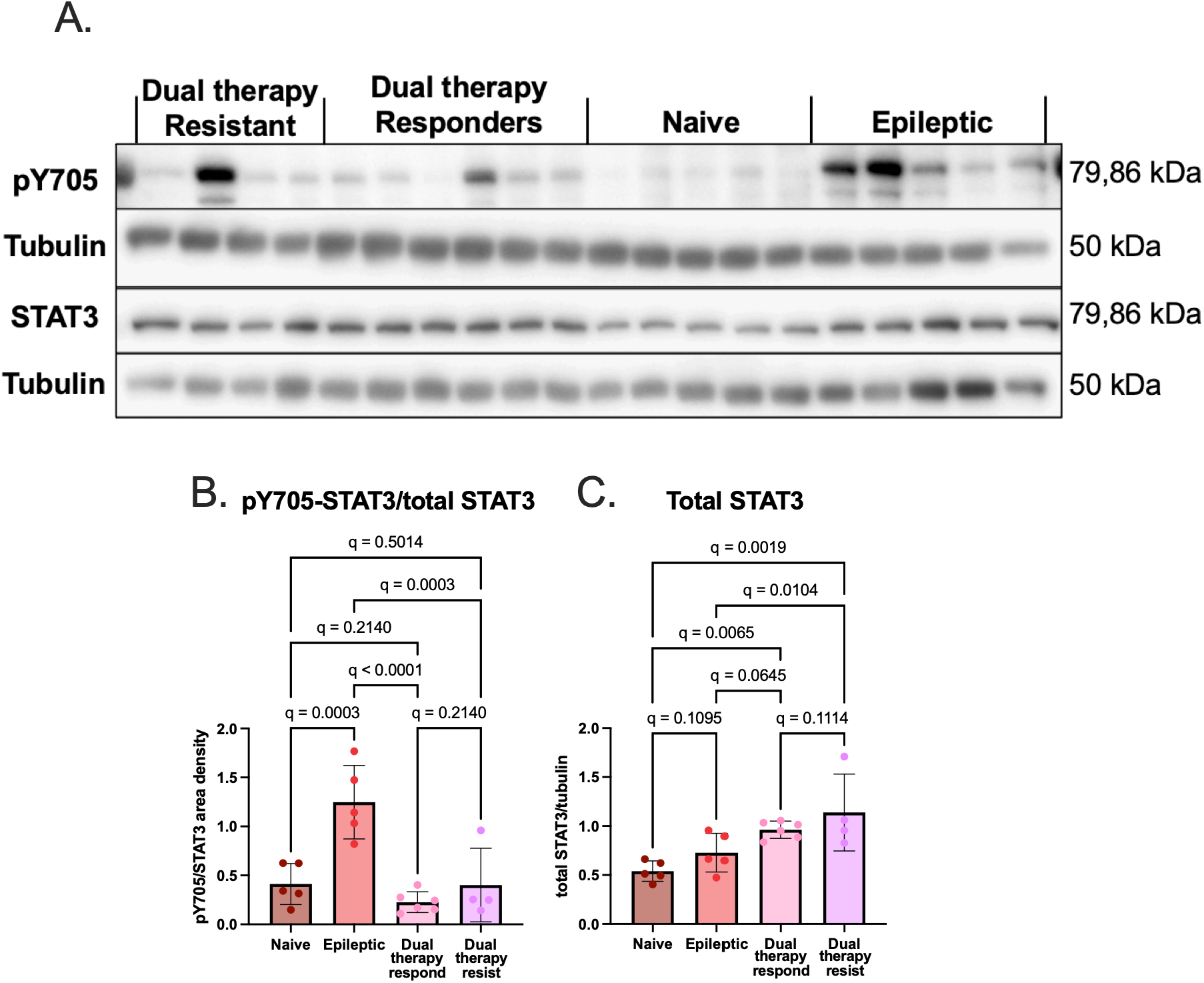
pY705-STAT3 is suppressed in CP690550 dual therapy resistant mice. **(A)** Representative western blots for pY705-STAT3 and STAT3 in the hippocampi of naïve, epileptic (20 weeks post SE), and dual therapy treated mice who (16 weeks post SE) who either responded or were resistant to treatment. Tubulin was used as a loading control. **(B)** Quantification of pY705-STAT3 and **(C)** total STAT3 in naïve, epileptic, and dual therapy treated mice who responded to were resistant to treatment. Statistical analysis performed using a One-way ANOVA with BKY post hoc.

## Discussion

Drug-resistant temporal lobe epilepsy remains one of the most difficult problems in clinical epileptology because the failure of an initial antiseizure medication strongly predicts a reduced likelihood of response to subsequent medications, even when those agents have distinct pharmacological mechanisms *(5–7)*. Here, we developed a preclinical add-on treatment paradigm in which chronically epileptic mice were first screened for response to carbamazepine (CBZ), a clinically established sodium-channel-targeting antiseizure medication, and then CBZ-resistant mice were treated with the JAK inhibitor tofacitinib/CP690550 while CBZ treatment was continued (dual therapy). We found that systemic kainic acid-induced epilepsy produced the expected heterogeneity of CBZ seizure-response, with most mice failing to achieve a clinically meaningful reduction in seizure frequency *(44–46, 48, 49)*. In this CBZ-resistant cohort, addition of CP690550 produced marked seizure suppression, with a majority of animals responding to treatment and many becoming free of behavioral seizures during the recording period. CP690550 also restored spatial working and short-term memory in mice for which CBZ alone was ineffective. Finally, we show that seizure-response for both CBZ and dual CP690550 treatment is uncoupled from cognitive-response arguing against the framework that effective seizure suppression will drive cognitive restoration.

A key strength of this study is that the treatment paradigm was anchored to pharmacokinetic validation. CBZ concentrations in brain declined over the 8-hour period following injection, and this decline was paralleled by loss of anticonvulsant efficacy in the fluoroethyl seizure threshold assay. The relationship between brain CBZ concentration and seizure threshold supports the interpretation that the CBZ dosing schedule used here provides biologically meaningful antiseizure exposure during behavioral monitoring. These findings are consistent with prior preclinical pharmacokinetic studies of prototype antiseizure medications*(50)* and provide an important internal control for interpreting CBZ nonresponse in chronically epileptic mice.

We also detected CP690550 in the brains of chronically epileptic mice with a brain-to-plasma ratio of approximately 0.12. This finding indicates that CP690550 has limited, but measurable, brain exposure in epileptic animals. The absence of detectable brain trospium chloride, a non-blood-brain-barrier-penetrant control compound *(51)*, supports the conclusion that CP690550 detection does not simply reflect contamination of the brain samples by residual blood. The limited central exposure is relevant for two reasons. First, the measured brain concentration exceeded 89% of reported IC50 values for JAK-1 heterodimers measured in an in vitro whole cell assay*(40)*, supporting the plausibility of direct target engagement in brain tissue. Second, the low brain-to-plasma ratio leaves open the possibility that the therapeutic effect of CP690550 is not mediated exclusively through actions in the central nervous system: peripheral immune cells, brain endothelial cells, vascular-associated cells, or other compartments at the blood-brain interface may contribute to seizure suppression and cognitive rescue. Together these data emphasizes that anti-epileptic therapy need not act only through classical neuronal antiseizure mechanisms.

The central translational finding is that CP690550 suppresses seizures in mice that remain refractory to CBZ. The International League Against Epilepsy defines drug-resistant epilepsy as failure of adequate trials of two tolerated and appropriately chosen antiseizure medications as measured via seizure-response *(6)*. Our model does not fully reproduce that clinical definition because animals were screened against a single antiseizure medication. However, failure of the first appropriately chosen medication is a clinically meaningful event: patients who do not respond to initial therapy have a substantially reduced probability of responding to the next antiseizure medication *(7)*, and early treatment failure is strongly associated with later pharmacoresistance in general *(5)*. Thus, CBZ resistance in this model should be interpreted as a preclinical approximation of an early pharmacoresistant state rather than as a complete model of clinical drug-resistant epilepsy. The effect of CP690550 were substantial: in animals classified as CBZ resistant, dual CP690550 treatment reduced both seizure frequency and total time spent seizing. Across the full treatment sequence, a large fraction (82%) of mice achieved at least a 50% reduction in seizure frequency. These data suggest that targeting JAK/STAT-linked inflammatory signaling can overcome seizure-resistance to a conventional antiseizure medication in a chronic acquired epilepsy model.

This result is particularly important because most current antiseizure medications primarily reduce neuronal excitability or synaptic transmission, whereas CP690550 targets a signal transduction pathway implicated in inflammatory and reactive cellular states. Drug resistance likely reflects multiple overlapping mechanisms, including altered drug transport, changes in drug targets, network reorganization, neuroinflammation, and disease severity *(4, 5, 52)*. The ability of CP690550 to suppress seizures after CBZ failure suggests that at least some component of pharmacoresistance in chronic temporal lobe epilepsy may be bypassed by targeting noncanonical, disease-associated signaling mechanisms rather than simply adding another conventional antiseizure medication. This does not imply that CP690550 is universally effective, because a subset of mice did not achieve seizure-response. Rather, the bimodal response suggests that JAK/STAT-dependent mechanisms may be dominant in one subgroup of chronically epileptic animals, whereas other mechanisms maintain seizures in another subgroup.

Among mice that were responsive to dual therapy, the magnitude of seizure suppression was striking. Most CP690550 seizure-responders became free of observed behavioral seizures during the recording window, and the reduction in seizure frequency was substantially greater than that seen in CBZ responders (100-fold vs 3-fold respectively).

However, while we acknowledge that the use of video-based seizure detection is an important limitation of this study, our previous work found close agreement between behavioral seizure suppression and EEG seizure suppression during CP690550 treatment *(24)*. Indeed, our previous work showed that after three days of CP690550 treatment the majority of mice were seizure free as measured by both 24/7 EEG and behavioral seizure recording*(24)*, justifying the use of video monitoring in this study.

The cognitive results extend the significance of CP690550 beyond seizure suppression. Cognitive impairment is common in temporal lobe epilepsy and often profoundly affects quality of life. Indeed, studies suggest that cognitive impairment may have a greater impact on quality of life than seizures themselves *(53, 54)*. In this study, CBZ failed to restore spatial working memory or short-term memory, whereas addition of CP690550 restored performance to levels comparable to naïve animals (Fig 4). These data are consistent with our previous finding that transient CP690550 treatment improves cognitive outcomes in chronic epilepsy *(24)*, and they suggest that JAK/STAT inhibition can rescue disease-associated cognitive dysfunction even in animals first exposed to an ineffective antiseizure medication. Importantly, cognitive restoration was not restricted to animals that exhibited seizure suppression. CBZ-treated mice remained cognitively impaired regardless of whether their seizure frequency improved, whereas CP690550 improved short-term memory even in mice that remained seizure non-responsive by the behavioral metrics used herein. This dissociation argues against a simple model in which cognition improves only because seizure frequency decreases.

The uncoupling of seizure and cognitive-response has several implications. First, it suggests that seizures and cognitive impairment may be parallel consequences of a shared disease process rather than a strictly linear cascade in which seizures alone drive cognitive decline. Second, it supports the idea that inflammatory JAK/STAT signaling may contribute directly to hippocampal dysfunction. Cognitive impairment has been correlated with seizure burden in many studies *(16, 18)*, but cognitive and behavioral deficits are also detectable near disease onset, before the first recorded seizure and before epilepsy onset or prolonged antiseizure medication exposure *(55, 56)*. These observations are compatible with a model in which epileptogenic injury, neuroinflammation, and circuit remodeling jointly produce both seizures and cognitive deficits. The ILAE definition of drug refractory epilepsy reflects seizure-response but does not incorporate a metric that address patients who continue to cognitively decline despite adequate seizure control. An example of this phenomenon is patients with late onset epilepsy who show well controlled seizures on existing ASMs but go on to face higher rates of dementia, cognitive impairment, stroke, and an up to 6-fold higher mortality rates compared to the general population*(57, 58)*.

The relevance of STAT3-linked inflammatory signaling to cognition may also extend beyond epilepsy, because JAK/STAT activation has been implicated in neuroinflammatory and cognitive phenotypes in models of Alzheimer’s disease and Huntington’s disease *(29–32, 59)*. The present data therefore support further investigation of JAK/STAT signaling as a modifiable pathway linking chronic brain inflammation to cognitive dysfunction.

Monitoring of STAT3 phosphorylation and activation provide mechanistic support for target engagement while also clarifying the limits of the current study. Consistent with our prior work *(24)*, chronically epileptic mice showed increased hippocampal pY705-STAT3. Two weeks of CBZ+CP690550 suppressed pY705-STAT3 in both seizure-responders and nonresponders. Thus, failure to suppress seizures in a subset of mice was not attributable to a gross failure of CP690550 to inhibit hippocampal STAT3 phosphorylation. This finding argues against a simple pharmacokinetic or target-access explanation for nonresponse and suggests that resistance to CP690550 may arise downstream of, parallel to, or independent from hippocampal STAT3 activation.

This study has some limitations. First, the model screens resistance to CBZ alone, whereas clinical drug-resistant epilepsy requires failure of two appropriate medications *(6, 24)*. The study therefore models an early or partial pharmacoresistant phenotype rather than the full clinical syndrome. However, it should be noted that failure to respond to a first appropriate ASM is highly predictive of drug resistance in general*(5, 7–9)*. Second, seizure outcomes were measured by behavioral video monitoring rather than continuous EEG. Though, we note that prior EEG and- video concordance during CP690550 treatment supports this approach *(24)*.

In summary, this study provides preclinical evidence that CP690550 can function as an effective add-on therapy in a chronic temporal lobe epilepsy after failure of CBZ. CP690550 suppressed seizures and restored spatial and short-term memory. These findings support a model in which inflammatory JAK/STAT signaling contributes to both seizure burden and cognitive dysfunction in chronic epilepsy, but in partially separable ways. More broadly and perhaps most importantly, the data suggest that pharmacoresistant epilepsy may be treatable by targeting disease-modifying inflammatory pathways rather than relying solely on additional antiseizure medications that modulate neuronal excitability.

