## Supplemental Figures for "Tofacitinib overrides first-line drug resistance in a pre-clinical model of epilepsy"

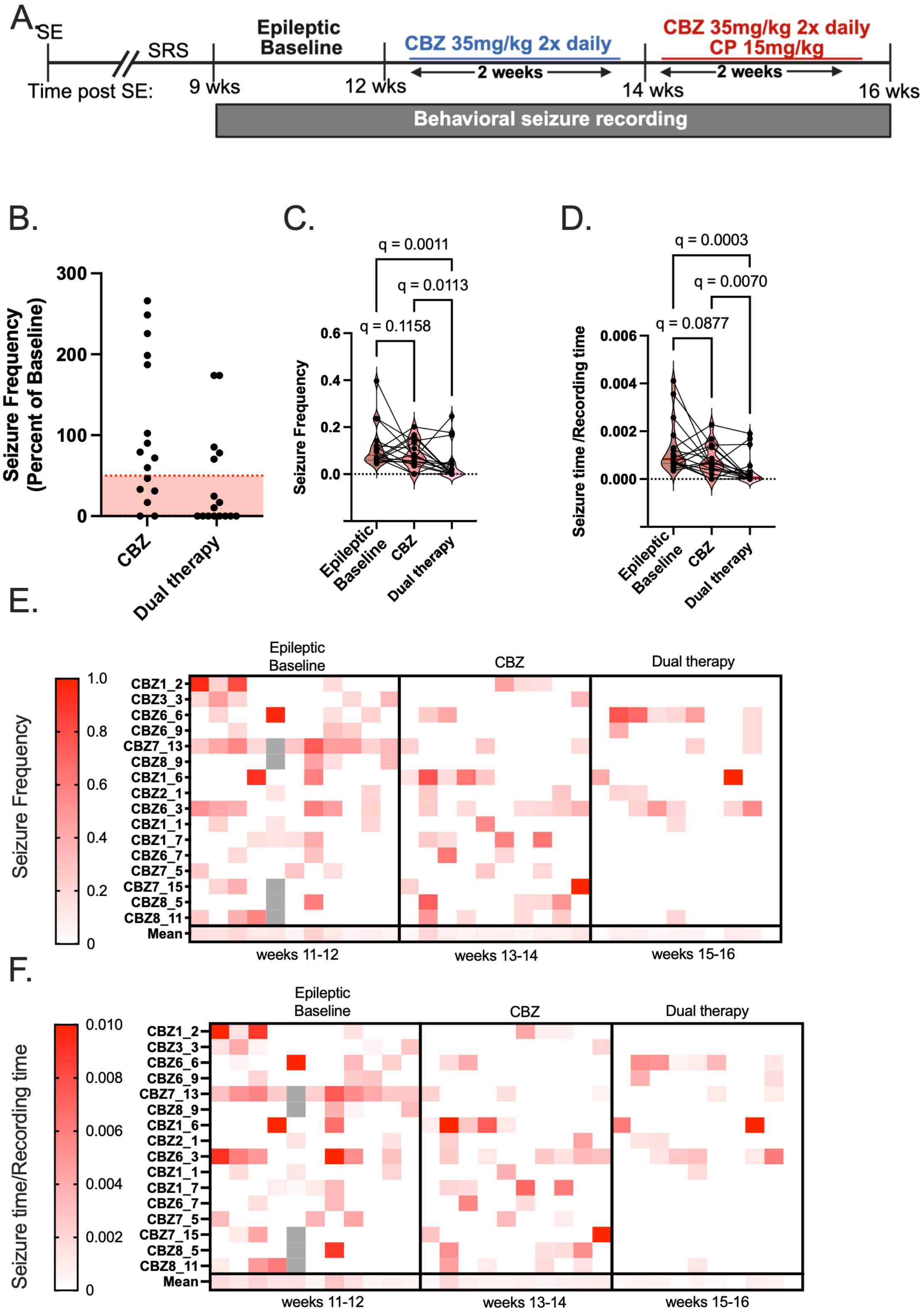


**Figure S1. CP690550/CBZ combination treatment suppresses seizures in chronic epilepsy. (A)** Experimental timeline. **(B)** Percent baseline seizure frequency, of CBZ and CBZ+CP690550 treated mice. A reduction in percent seizure frequency of 50% or more, the shaded red area, indicates a mouse responded to treatment. Each point represents a mouse. Violin plots depict the **(C)** seizure frequency and **(D)** time spent seizing for baseline, CBZ treated, and CBZ+CP690550 treated mice. The median is represented by a bold maroon line. Statistics were calculated using the Friedman test with BKY post hoc. Heatmaps depict the changes in **(E)** seizure frequency and **(F)** time spent seizing for individual mice across the treatment paradigm.

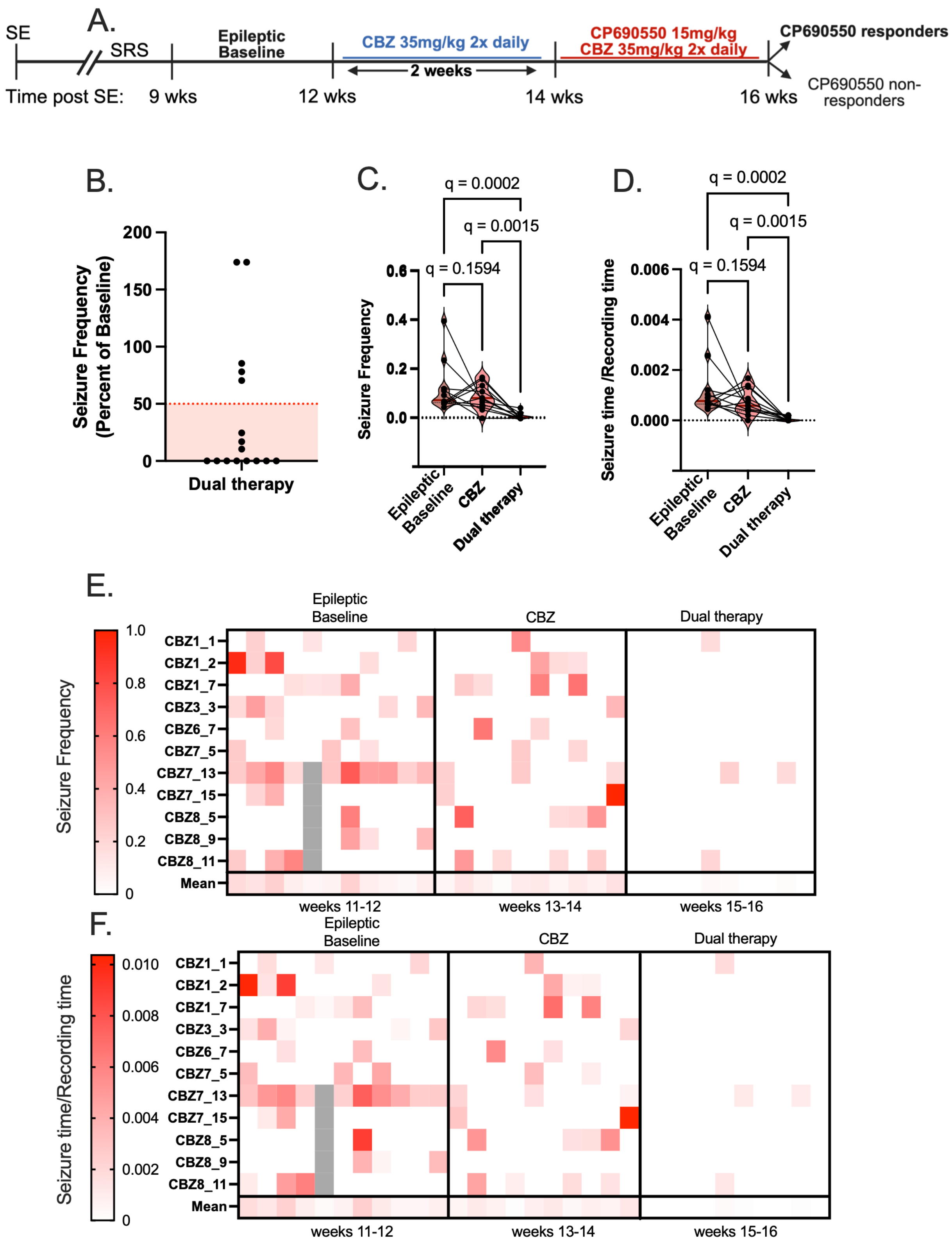

A.
SE
SRS
Epileptic Baseline
CBZ 35mg/kg 2x daily
CP690550 15mg/kg
CBZ 35mg/kg 2x daily
CP690550 responders
2 weeks
Time post SE:
9 wks
12 wks
14 wks
16 wks
CP690550 non-responders
B.
Seizure Frequency (Percent of Baseline)
200
150
100
50
0
Dual therapy
C.
q = 0.0002
q = 0.0015
q = 0.1594
Seizure Frequency
0.6
0.4
0.2
0.0
Epileptic Baseline
CBZ
Dual therapy
D.
q = 0.0002
q = 0.0015
q = 0.1594
Seizure time /Recording time
0.006
0.004
0.002
0.000
Epileptic Baseline
CBZ
Dual therapy
E.
Seizure Frequency
1.0
0.8
0.6
0.4
0.2
0
Epileptic Baseline
CBZ
Dual therapy
CBZ1_1
CBZ1_2
CBZ1_7
CBZ3_3
CBZ6_7
CBZ7_5
CBZ7_13
CBZ7_15
CBZ8_5
CBZ8_9
CBZ8_11
Mean
weeks 11-12
weeks 13-14
weeks 15-16
F.
Seizure time/Recording time
0.010
0.008
0.006
0.004
0.002
0
Epileptic Baseline
CBZ
Dual therapy
CBZ1_1
CBZ1_2
CBZ1_7
CBZ3_3
CBZ6_7
CBZ7_5
CBZ7_13
CBZ7_15
CBZ8_5
CBZ8_9
CBZ8_11
Mean
weeks 11-12
weeks 13-14
weeks 15-16

**Figure S2. CP690550/CBZ combination treatment profoundly suppresses seizures in CP690550 responders (A)** Experimental timeline. **(B)** Percent baseline seizure frequency, of CBZ+CP690550 treated mice. A reduction in percent seizure frequency of 50% or more, the shaded red area, indicates a mouse responded to treatment, only CBZ+CP690550 responders were included in the rest of the analysis. Each point represents a mouse. Violin plots depict the **(C)** seizure frequency and **(D)** time spent seizing for baseline, CBZ treated, and CBZ+CP690550 treated mice. The median is depicted as a bold maroon line. Statistics were calculated using the Friedman test with BKY post hoc. Heatmaps depict the changes in **(E)** seizure frequency and **(F)** time spent seizing for individual mice across the treatment paradigm.

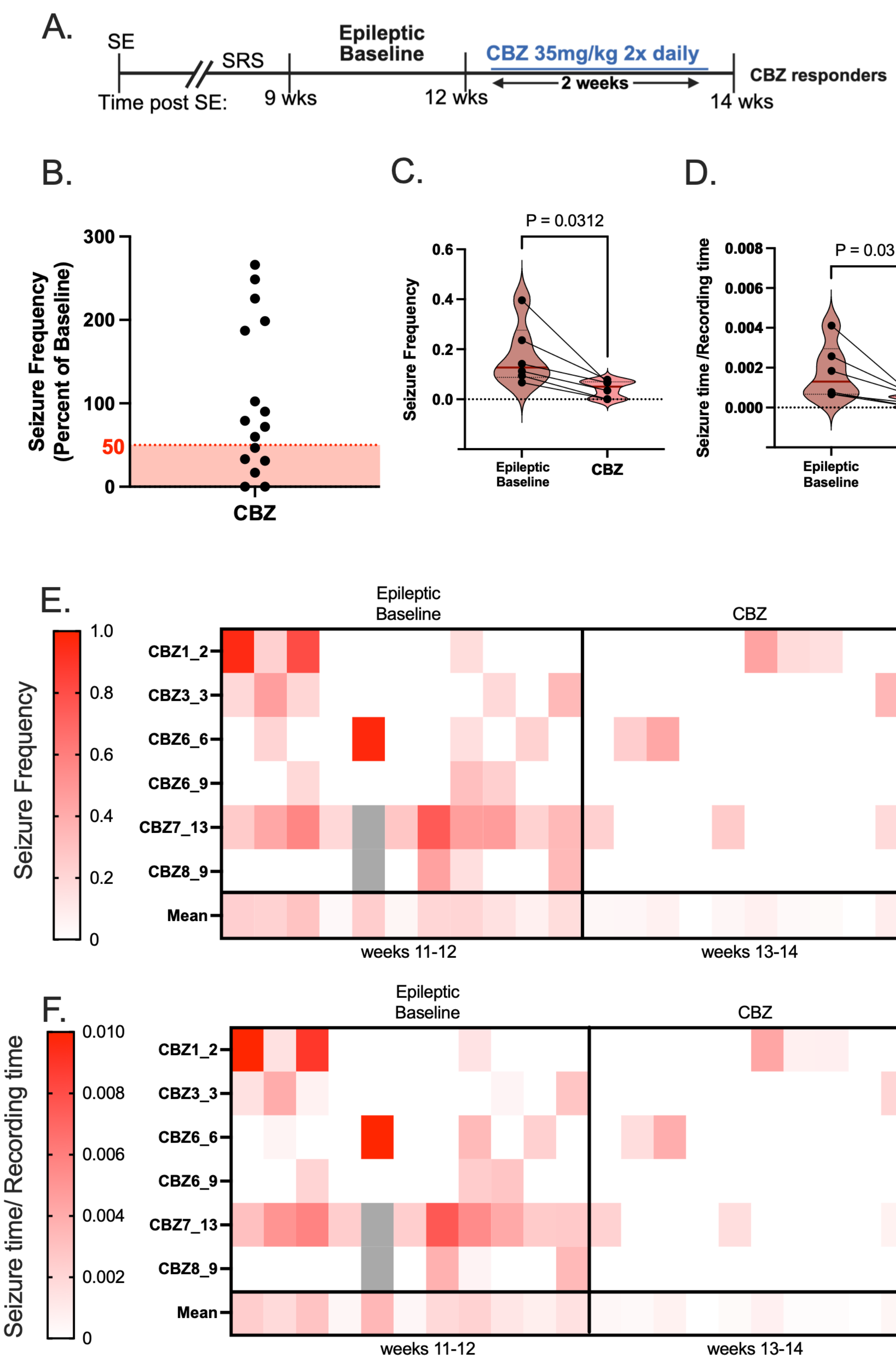


**Figure S3. CBZ responders show moderate seizure suppression (A)** Experimental timeline **(B)** Percent baseline seizure frequency, of CBZ treated mice. A reduction in percent seizure frequency of 50% or more, the shaded red area, indicates a mouse responded to treatment, only CBZ responders were included in the rest of the analysis. Each point represents a mouse. Violin plots depict the **(C)** seizure frequency and **(D)** time spent seizing for baseline, and CBZ treated mice. The median is depicted as a bold maroon line. Statistics were calculated using the Wilcoxon matched pairs signed rank test. Heatmaps depict the changes in **(E)** seizure frequency and **(F)** time spent seizing for individual mice across the treatment paradigm.

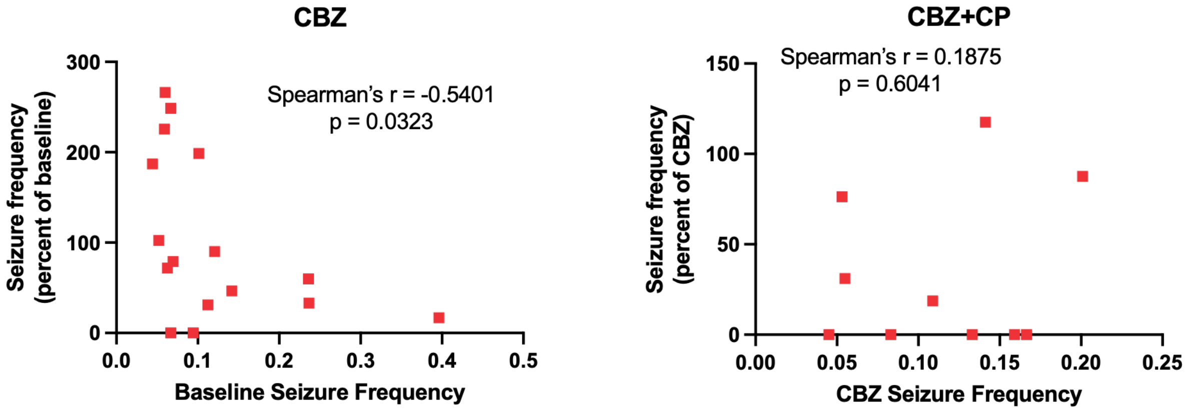


**Figure S4. Seizure severity is not sufficient to explain drug resistance.**
**(A)** Correlation analysis shows slight significant negative correlation (r = -0.541, p = 0.0323) between baseline seizure frequency and percent seizure reduction on CBZ treatment. **(B)** We see no correlation between seizure frequency and percent seizure reduction on CBZ+CP690550 in CBZ resistant mice (r = 0.1875, p = 0.6041). Analysis done using Spearman's correlation.

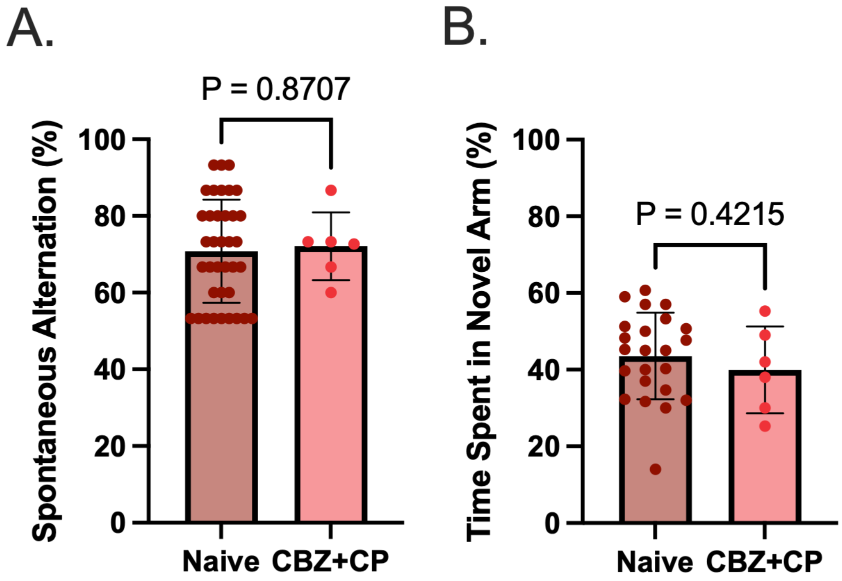


**Figure S5. CBZ+CP690550 combination treatment does not impair naïve animals**. Bar graphs depict analysis of **(A)** spontaneous alternations and **(B)** time spent in novel arm for naïve mice and naïve mice treated with CBZ and CP690550 combination treatment. Each point represents a mouse. Statistical analysis performed using Wilcoxon Ranked Sum test.

Seizure frequency during experiment

| Cohort ID | Treatment | BASELINE MEAN | CBZ | CP +CBZ MEAN |
|---|---|---|---|---|
| CBZ1_1 | CBZ then CBZ+CP | 0.06946747 | 0.05496183 | 0.01706161 |
| CBZ1_2 | CBZ then CBZ+CP | 0.23655215 | 0.07835524 | 0 |
| CBZ1_6 | CBZ then CBZ+CP | 0.10118189 | 0.20096785 | 0.17589513 |
| CBZ1_7 | CBZ then CBZ+CP | 0.06690987 | 0.16643367 | 0 |
| CBZ2_1 | CBZ then CBZ+CP | 0.05196208 | 0.05314489 | 0.04054337 |
| CBZ3_3 | CBZ then CBZ+CP | 0.11243397 | 0.03485839 | 0 |
| CBZ6_3 | CBZ then CBZ+CP | 0.23604707 | 0.14123884 | 0.16605805 |
| CBZ6_6 | CBZ then CBZ+CP | 0.14172966 | 0.06623854 | 0.24656053 |
| CBZ6_7 | CBZ then CBZ+CP | 0.04441725 | 0.08306744 | 0 |
| CBZ6_9 | CBZ then CBZ+CP | 0.0668275 | 0 | 0.05712075 |
| CBZ7_5 | CBZ then CBZ+CP | 0.062641 | 0.04504594 | 0 |
| CBZ7_13 | CBZ then CBZ+CP | 0.39620694 | 0.06716882 | 0.04155296 |
| CBZ7_15 | CBZ then CBZ+CP | 0.05900315 | 0.13310896 | 0 |
| CBZ8_5 | CBZ then CBZ+CP | 0.05978577 | 0.15916341 | 0 |
| CBZ8_9 | CBZ then CBZ+CP | 0.09422489 | 0 | 0 |
| CBZ8_11 | CBZ then CBZ+CP | 0.1207766 | 0.10879991 | 0.02034818 |
| | **Means** | **0.12001045** | **0.8703461** | **0.04782129** |

**Table S1. Seizure frequency for all mice during experiment.** Seizure frequency was measured in seizures per hour.

**Time seizing during experiment**

| Cohort ID | Treatment | BASELINE MEAN | CBZ | CP+CBZ |
|---|---|---|---|---|
| CBZ1_1 | CBZ then CBZ+CP | 0.00068564 | 0.00037405 | 0.00018009 |
| CBZ1_2 | CBZ then CBZ+CP | 0.00257333 | 0.00058128 | 0 |
| CBZ1_6 | CBZ then CBZ+CP | 0.00130011 | 0.00227544 | 0.00190382 |
| CBZ1_7 | CBZ then CBZ+CP | 0.00065122 | 0.00167744 | 0 |
| CBZ2_1 | CBZ then CBZ+CP | 0.00035148 | 0.00067621 | 0.00029055 |
| CBZ3_3 | CBZ then CBZ+CP | 0.00066385 | 0.00020614 | 0 |
| CBZ6_3 | CBZ then CBZ+CP | 0.003557 | 0.00142628 | 0.00144747 |
| CBZ6_6 | CBZ then CBZ+CP | 0.00183602 | 0.00055936 | 0.00168849 |
| CBZ6_7 | CBZ then CBZ+CP | 0.00045042 | 0.00073899 | 0 |
| CBZ6_9 | CBZ then CBZ+CP | 0.00067422 | 0 | 0.0005556 |
| CBZ7_5 | CBZ then CBZ+CP | 0.00102965 | 0.00042466 | 0 |
| CBZ7_13 | CBZ then CBZ+CP | 0.0041115 | 0.00044944 | 0.00021158 |
| CBZ7_15 | CBZ then CBZ+CP | 0.00053273 | 0.00132829 | 0 |
| CBZ8_5 | CBZ then CBZ+CP | 0.00090094 | 0.0013854 | 0 |
| CBZ8_9 | CBZ then CBZ+CP | 0.00077262 | 0 | 0 |
| CBZ8_11 | CBZ then CBZ+CP | 0.00120126 | 0.00086588 | 0.00012435 |
| | **Means** | **0.00133075** | **0.00081055** | **0.00040012** |

**Table S2. Time seizing for all mice during experiment.** Time seizing was measured as seconds seizing over total recording time (sec).
